# The dynamics of ERK signaling in melanoma, and the response to BRAF or MEK inhibition, are cell cycle dependent

**DOI:** 10.1101/306571

**Authors:** Chloe M. Simpson, Nicola Ferrari, Fernando Calvo, Chris Bakal

## Abstract

Activating BRAF mutations drive melanoma tumorigenesis and metastasis by constitutively activating MEK and ERK, and small molecule inhibitors (SMIs) of BRAF or MEK have shown promise as melanoma therapeutics. However, the development of resistance to these inhibitors in both the short- and longterm is common; warranting investigation into how these SMIs influence ERK signaling dynamics. By quantitative single cell imaging of an ERK activity reporter in living cells, we describe both intra- and inter-cell heterogeneity in ERK activity in isogenic melanoma populations harboring a BRAFV600E mutation. This heterogeneity is largely due to a cell-cycle dependent bifurcation in ERK activation. Moreover, we show there are cell-cycle dependent responses in ERK activity following BRAF or MEK inhibition. Prior to, but not following, CDK4/6-mediated passage through the Restriction Point (RP) ERK activity is sensitive to BRAF and MEK inhibitors. In contrast, in cells that have passed the RP, ERK activity will remain elevated even in the presence of BRAF or MEK inhibition until mitosis. We propose that ERK activity in the presence of activating BRAF mutations is regulated by both positive and negative feedback loops that are engaged in cell-cycle dependent fashions. CDK4/6 inhibition sensitizes ERK activity to BRAF or MEK inhibition by preventing passage the transition from a BRAF/MEK dependent to independent state. Our results have implications for the use of MEK and BRAF inhibitors as melanoma therapeutics, and offer a rational basis for the use of these inhibitors in combination with CDK4/6 inhibition during cancer therapy.

## Introduction

Extracellular signal-regulated kinase (ERK) regulates a diverse array of distinct, and sometimes even opposing, behaviors (1). ERK activity is frequently upregulated in cancers following activating mutations in RAS (*NRAS* and *KRAS*) and *BRAF* genes,(2,3), and/or mutations in negative regulators that inhibit the activity of ERK such as NF1, c-KIT or RASA2 (4, 5, 6, 7). Understanding how ERK activity regulates cell fate in cancer cells is key to developing and deploying therapies that suppress ERK activation. Such insight is particularly relevant to melanomas, which are almost all dependent on ERK activation for their progression, but where promising inhibitors of the ERK pathway appear to have limited effectiveness clinically (8, 9, 10, 11, 12, 13, 14, 15).

In normal cells, there is cell-to-cell variability in the dynamics of ERK activation in response to the same stimulus (16). Such cell-to-cell variation likely contributes to different outcomes during cell fate decisions (23,24). These variations can be a consequence of noise in the activity of upstream activators such as MEK (18), or the duration of the activating signal (i.e. pulsatile versus transient) (19). Additional factors such as the spatial position of cells within a multi-cellular environment (16), differences in protein abundance (20), and/or cell shape (21,22) could potentially also underpin cell-to-cell variability in ERK activity.

While in some populations ERK activation appears to be an “all-or-none” event (25,26,27), ERK activity has also been demonstrated to oscillate in both frequency-independent (28), and frequency-modulated (29) fashions; as well as having dampened oscillatory activity (**Error! Reference source not found.**, 30). Thus, some normal cells also display within-cell variability in ERK dynamics. Most likely, the dynamics of ERK activation are influenced by similar factors to those which have been observed to affect cell-to-cell variation, as well as cell- and cue-specific signaling network topologies that both activate ERK, and modulate its activity through positive and negative feedback (32,33,25,34,36,37).

Most of our understanding of ERK activation dynamics comes from normal cell models, where ERK activity is tightly modulated, and the full dynamic range of ERK activity in response to a cue can be explored. But how activating mutations in upstream kinases such as BRAF affect cell-to-cell variation in ERK activity and response in cancer cells is less clear. Furthermore, little is understood as to how ERK activity responds in single cells with activating mutations in BRAF or MEK following chemical inhibition of these kinases. In the absence of such studies, we have little insight into how therapies that are given to cancer patients affect ERK signaling dynamics in individual cells – which could dictate the timing and dose of therapy (39), and be important in the emergence of therapeutically resistant sub-populations (19).

### ERKTR ratio illustrates cellular heterogeneity in ERK activity

To quantify the dynamics of ERK activity in single cells immediately following the exposure of cells to commonly used small-molecule inhibitors of RTK-RAS-ERK signaling in BRAF mutant melanoma cells, we engineered a human A375p melanoma cell line to constitutively express the <u>E</u>RK translocation <u>r</u>eporter (ERKTR) (40, 41). A375p^ERKTR^ cells harbor the BRAFV600E mutation and were derived from a primary melanoma site (42, 43, Supplementary Figure 1). ERKTR is a fluorescently tagged single peptide reporter that contains both an ERK docking site, and consensus motifs for ERK phosphorylation (Figure 1A, 40). In an un-phosphorylated state, ERKTR is localized to the nucleus due to the presence of NLS-sites in the peptide (Figure 1B). When ERK activity increases, ERKTR is phosphorylated by ERK, which exposes cryptic NES sites, and ERKTR is exported to the cytoplasm (Figure 1C).

That ERKTR is a specific readout for ERK activity has been well established. Whilst the phosphorylation sites on ERKTR are not specific for ERK (44), the presence of a specific ERK docking site on ERKTR ensures that other proline-directed kinases do not phosphorylate ERKTR(40). The ERK docking site is comprised of ERK binding domains found in ELK1 including a D-domain (<u>K</u>G<u>RK</u>PRD<u>LEL</u>), and an FXFP motif (45, 46, 47). Though both JNK and ERK can bind the ELK1 D-domain (48), only ERK binds the FXFP motif (49). Thus, these two ERK binding sites act in an additive fashion to dictate the specificity of ERKTR (46).

**Figure 1.**
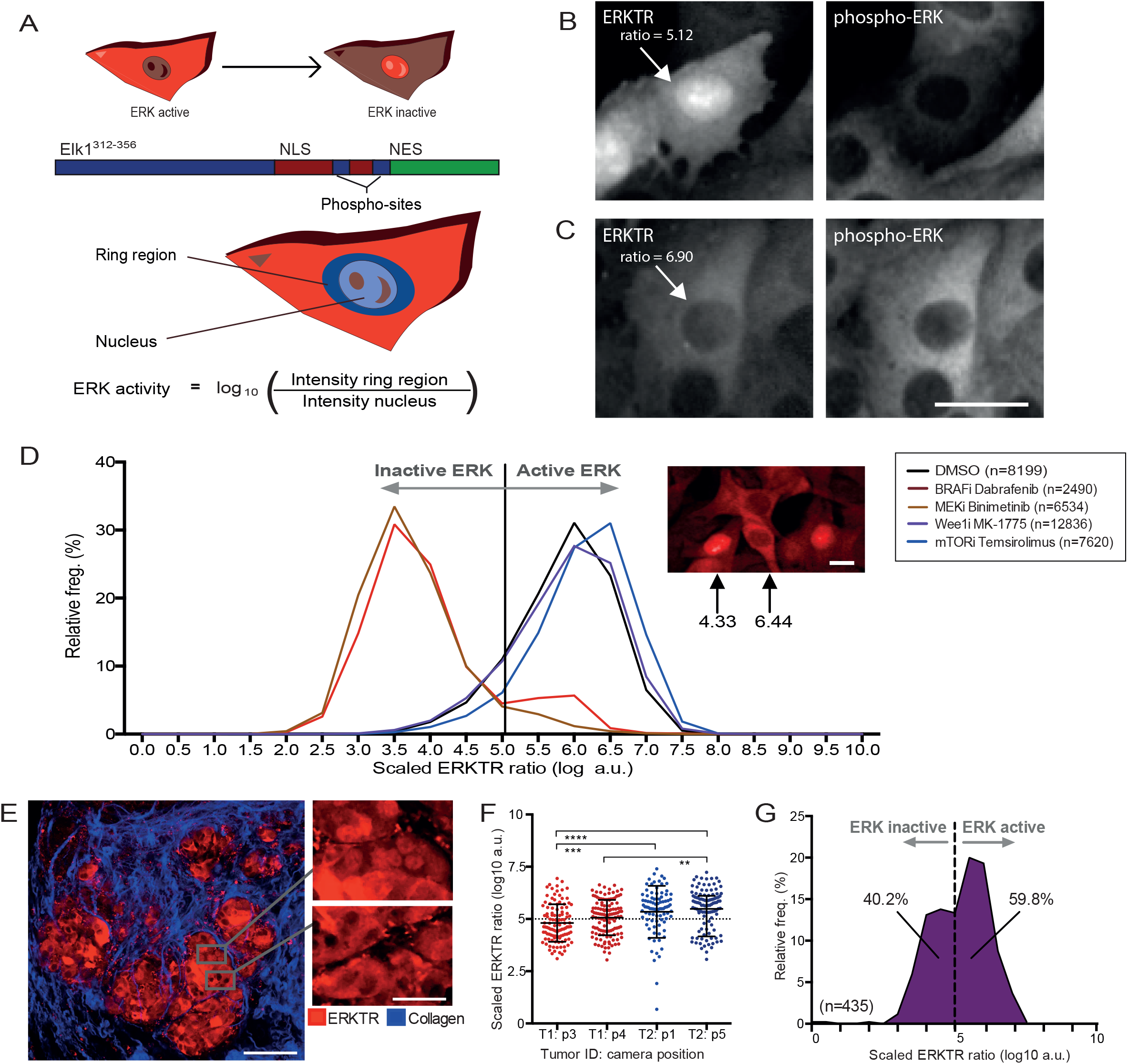
Cell-to-cell variability in ERK activation in BRAFV600E melanoma cells. (A) ERKTR is a reporter of ERK activity (Regot et al, 2014). (B) Phospho-ERK staining in A375p^ERKTR^ where ERK is inactive. (C) Phospho-ERK staining in A375p^ERKTR^ where ERK is active; scale bar = 25μm. (D) Frequency distribution of ERKTR ratio following treatment by: BRAF inhibitor (Dabrafenib), MEK inhibitor (Binimetinib), Wee1 inhibitor (MK-1775) and mTOR inhibitor (Temsirolimus) for 24hours; scale bar = 25μm (E) In vivo ERK activity (red) in single melanoma cells visualized by multiphoton intravital microscopy of subcutaneous tumours. In blue, collagen fibers obtained by second harmonic generation. Scale bars, 300μm (left panel) and 45μm (right panel). (F) Quantification of ERKTR ration from tumor xenografts. From left: tumor 1: camera position 3 (T1:p3), tumor 1: camera position 4 (T1:p4), tumor 2: camera position 1 (T2:p1), and tumor 2: camera position 5 (T2:p5). (G) Histogram of ERKTR ratios from compiled tumor images.

Confirming that ERKTR is specific reporter for ERK, and not other MAPKs such as JNK or p38, Regot et al., demonstrated that JNK or p38 inhibitors to not affect ERKTR translocation dynamics following stimulation by bFGF2 (40). Furthermore, ERKTR has been used to monitor ERK activity in parallel with an another ERK activity - EKAR FRET based ERK reporter (50), and the activity of both is largely identical (51). In addition, a form of ERKTR has been generated to monitor ERK activity during *in vivo* in *C. elegans*, and genetic perturbations, that are predicted to downregulate (*mpk1* null mutant) or upregulate (Lin-45/BRAF expression) ERK activity result in predicted effects on ERK activity (52). Taken together these data demonstrate that ERKTR is a specific reporter of ERK activation.

Here we aimed to assess the impact of a BRAFV600E mutational background on the dynamics of ERK signaling and how those dynamics are altered in response to clinically relevant inhibitors targeting the ERK pathway.

## Results

To validate the ability of ERKTR to serve as a readout for the BRAF-MEK-ERK signaling pathway in single melanoma cells, we fixed: A375p^ERKTR^ cells grown in normal conditions (10% serum); A375p^ERKTR^ cells treated with the BRAF inhibitor Dabrafenib; or A375p^ERKTR^ cells treated with the MEK inhibitor Binimetinib for 24 hours. Fixed cells were labeled with anti-phosphoERK antibody for the Tyr202/Thr204 MEK phosphorylation site. BRAF-MEK-ERK activity is determined by quantifying the log10 of cytoplasmic:nuclear ratio of ERKTR intensity, or single phospho-ERK intensity, rescaled to a 0-10 range (Figure 1A; Methods). Rescaling was done taking the minimum and maximum across the whole data set, preserving baseline values. Phospho-ERK levels and the ratio of cytoplasmic:nuclear ERKTR (hereafter referred to as ‘ERKTR ratio’) were positively correlated across conditions (Supplementary Figure 2a). Thus ERKTR localization is a readout of ERK phosphorylation on its activation sites.

Due to the implications of our study we also sought to validate that the ERKTR was not cross-reactive with CDK activity. In order to show this, we treated A375p^ERKTR^ cells with CDK inhibitors: CDK4/6 inhibitor Palbociclib (100uM), CDKi inhibitor Flavopiridol (100uM) and CDK1 inhibitor R03306 (100uM) gave no significant change in ERKTR ratio at 24hours. CDK4/6i gave a small, but significant change in ERKTR ratio at 2hours, mean difference: 0.169 (Supplementary Figure 2C). This confirms that ERKTR has little detectable cross-reactivity with CDK inhibitors.

The decrease in both phosphoERK levels and ERKTR ratio following BRAF or MEK inhibition, demonstrates that the ERKTR ratio faithfully responds to inhibition of BRAF-MEK-ERK activity in single melanoma cells. Notably however, there is considerable heterogeneity in both phospho-ERK and ERKTR ratios in melanoma cells after treatment with BRAF or MEK inhibitors. To characterize the heterogeneity of BRAF-MEK-ERK pathway activity, we plotted histograms of ERKTR ratios in single cells from untreated populations, and those treated with small molecule inhibitors (SMIs) of BRAF-MEK-ERK signaling. As negative controls we quantified ERK activity in A375p^ERKTR^ cells treated with Wee1i (MK-1775) and mTORi (Temsirolimus) inhibitors (Figure 1D, Supplementary Figure 3). In unperturbed cells (black curve, Figure 1D), the mean ERKTR ratio is 5.86, indicating that most cells have cytoplasmic ERKTR - and active BRAF-MEK-ERK signaling. However, the distribution of ERKTR ratios in normal cell populations exhibited a considerable negative skew (skewness=-0.629); thus, in normal A375p^ERKTR^ there is also significant sub-population of cells with low BRAF-MEK-ERK activity (Figure 1D). Conversely, whilst the mean ERKTR ratio is significantly decreased on a population level in A375p^ERKTR^ melanoma cells treated with either BRAF or MEK inhibitors compared to control cells, we observed that even in the presence of these inhibitors a fraction of A375p^ERKTR^ cells exhibited high ERKTR ratios that are comparable to those observed in control treated cells. These data suggest that even though the BRAF kinase is constitutively active in A375p cells, this constitutive activity does not translate to constitutive BRAF-MEK-ERK activity in all cells. Furthermore, our data shows that subpopulations of BRAFV600E expressing cells maintain high levels of BRAF-MEK-ERK activity, at least briefly, when exposed to BRAF or MEK inhibitors that effectively suppress ERK activity in other sub-populations.

To determine if melanoma cells exhibit similar heterogeneity in BRAF-MEK-ERK activity *in vivo*, we generated tumors of A375p^ERKTR^ cells in two immune-compromised mice, and imaged each tumor in live mice using two-photon confocal microscopy (Methods). We observed considerable heterogeneity in ERKTR ratios *in vivo*, including regional variation with “pockets” of cells where BRAF-MEK-ERK activity was decreased (Figure 1E). There was also significant variation in ERKTR ratios between the two tumors (Figure 1F). When we compiled data from both tumors, we observed that a significant population of cells (40.2%) had inactive BRAF-MEK-ERK pathway activity (Figure 1G). Thus, the existence of an activating BRAFV600E mutation in melanoma cells is insufficient to maintain constitutively high levels of BRAF-MEK-ERK activity *in vivo*.

### ERK activity bifurcates after mitosis

To determine the basis for population-level heterogeneity in BRAF-MEK-ERK activity, we quantified ERKTR ratios in A375p^ERKTR^ cells over time. ERK activity has been observed to exhibit fluctuations in other cell types during the cell cycle (29, 52, 53, 54), therefore, we engineered A375p^ERKTR^ cells to also express a fluorescently tagged version of Proliferating Cell Nuclear Antigen (PCNA) (Figure 2A) – which allows simultaneously quantification ERKTR ratios during different cell cycle phases. To demark S-phase we used automated image analysis methods (58) to identify a characteristic pattern of PCNA intensity increase, coinciding with a sharp peak in PCNA foci levels (56, Methods). From the timing of S-phase and mitosis, G1 and G2 phases can subsequently be inferred. After image analysis, we generated “tracks” of PCNA and ERKTR features over time, which can be aligned to different cell cycle transitions *in silico*.

**Figure 2.**
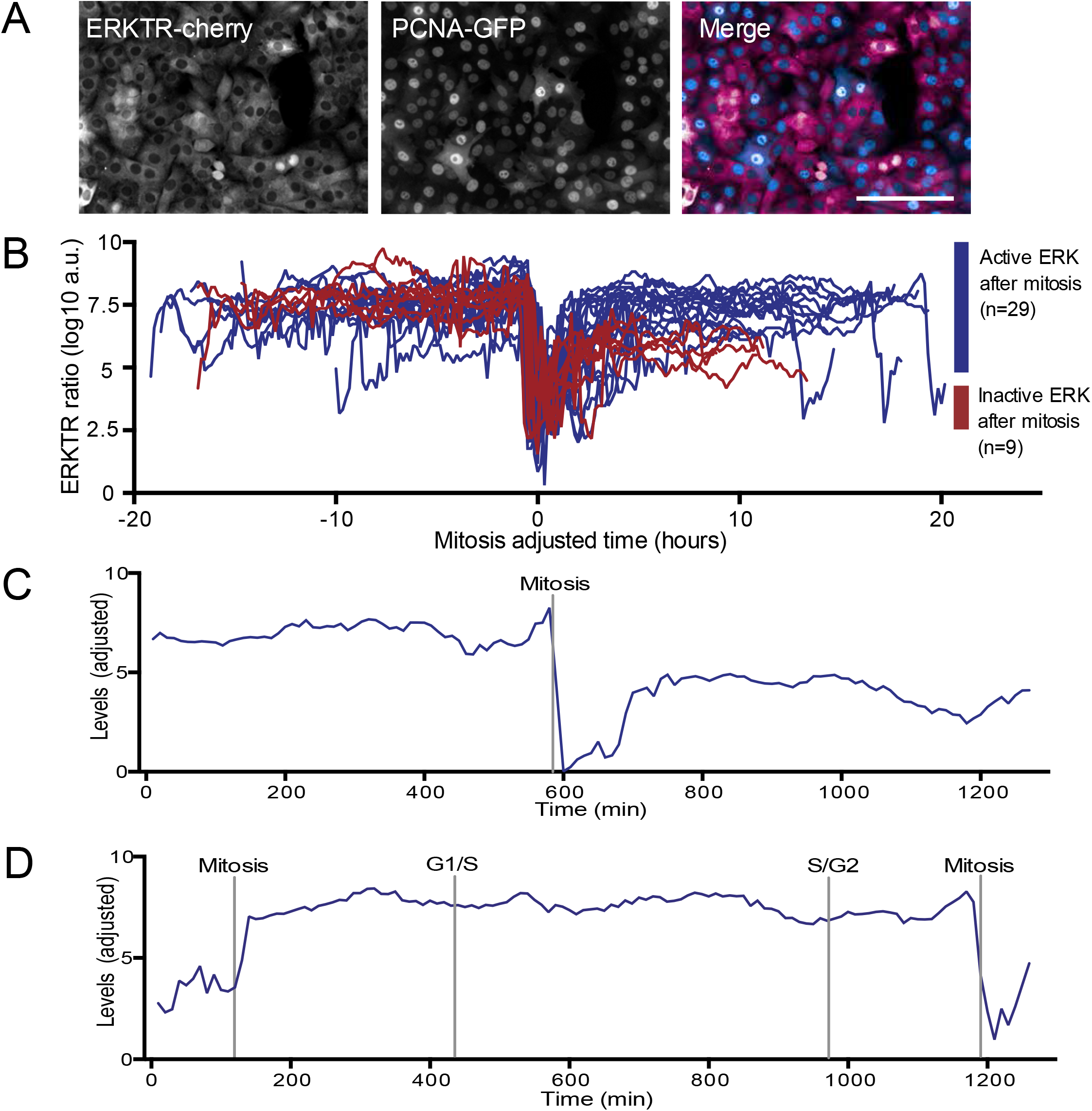
Bifurcation of ERK activity during the cell cycle. (A) Images of ERKTR and PCNA in labeled A375p^ERKTR^ cells. Scale bar = 200μm. (B) ERKTR ratio was quantified over time in single A375p^ERKTR/PCNA^ cells (DMSO treated), and is described by a single track. All tracks are aligned to mitosis. Due to nuclear envelope breakdown during mitosis, ERKTR ratios cannot be measured at mitosis, resulting in a dip in ratios. There is a bifurcation in ERK activity at mitosis that occurs in G1. 9 of 38 cells (23%) switched ERK off in G1, while 29 of 38 cells (77%) retained active ERK. (C) ERKTR ratio in a single cell where the ERKTR ratio drops in activity goes in G1 and the cells enters G1 arrest. D) ERKTR ratio in a proliferating cell.

Over multiple cell cycles, there was considerable intra-cell variation in ERKTR ratios. Specifically, in ~23% of cells, ERKTR ratios in daughter cells (D) decreased by as much as 5-fold compared to their mother cells (M) upon mitotic exit (Figure 2B). The ERKTR ratios in a representative cell whose ERKTR ratio falls upon G1 arrest are shown in Figure 2C, and a representative cell maintaining high ERKTR ratio throughout the cell cycle is shown in Figure 2D. We did not observe large fluctuations in ERKTR ratios outside the transition between mitosis and G1 in A375p cells, but observed oscillatory ERK dynamics in hTERT-RPE epithelial cells (Supplementary Figure 4A,B.). We also ruled out the existence of high frequency fluctuations in ERKTR ratios by imaging live cells at 1 min intervals (Supplementary Figure 4C). Thus, A375p melanoma cells exhibited considerable intra-cell variability in BRAF-MEK-ERK pathway activity, which can at least in part explain cell-to-cell differences observed at the population level.

### The response of ERKTR to BRAF and MEK inhibitors is dependent on cell cycle stage

We next wanted to determine how the BRAF (Dabrafenib, Vemurafenib), MEK (Binimetinib, Selumetinib), ERK (SCH772984) and RAS/Farnesyl Transferase (Lonafarnib) inhibitors, affected ERK dynamics over the cell cycle. Therefore, we imaged A375p^ERKTR/PCNA^ cells for 48 hours following the addition of SMIs. PCNA and ERKTR traces generated by imaging a single cell in Dabrafenib are shown in supplementary figure 4D,E. As negative controls, we monitored ERK dynamics following exposure of cells to Wee1 inhibitor or mTORi inhibitor (Temsirolimus) to show that any phenotype was due to the effect of inhibiting the BRAF-MEK-ERK pathway rather than a general cytotoxic effect of SMIs. To show how BRAF-MEK-ERK activity changes over time we aligned individual cell tracks to mitotic exit, or the first mitotic exit if the cell divided twice (56). Cells not observed to divide over the time course have been aligned with the last tracked value at -1h before mitosis, demarked with a double line (Figure 3). Each row represents a single cell track with a color scale normalized to the ERK ratio across the whole data set (Methods).

**Figure 3.**
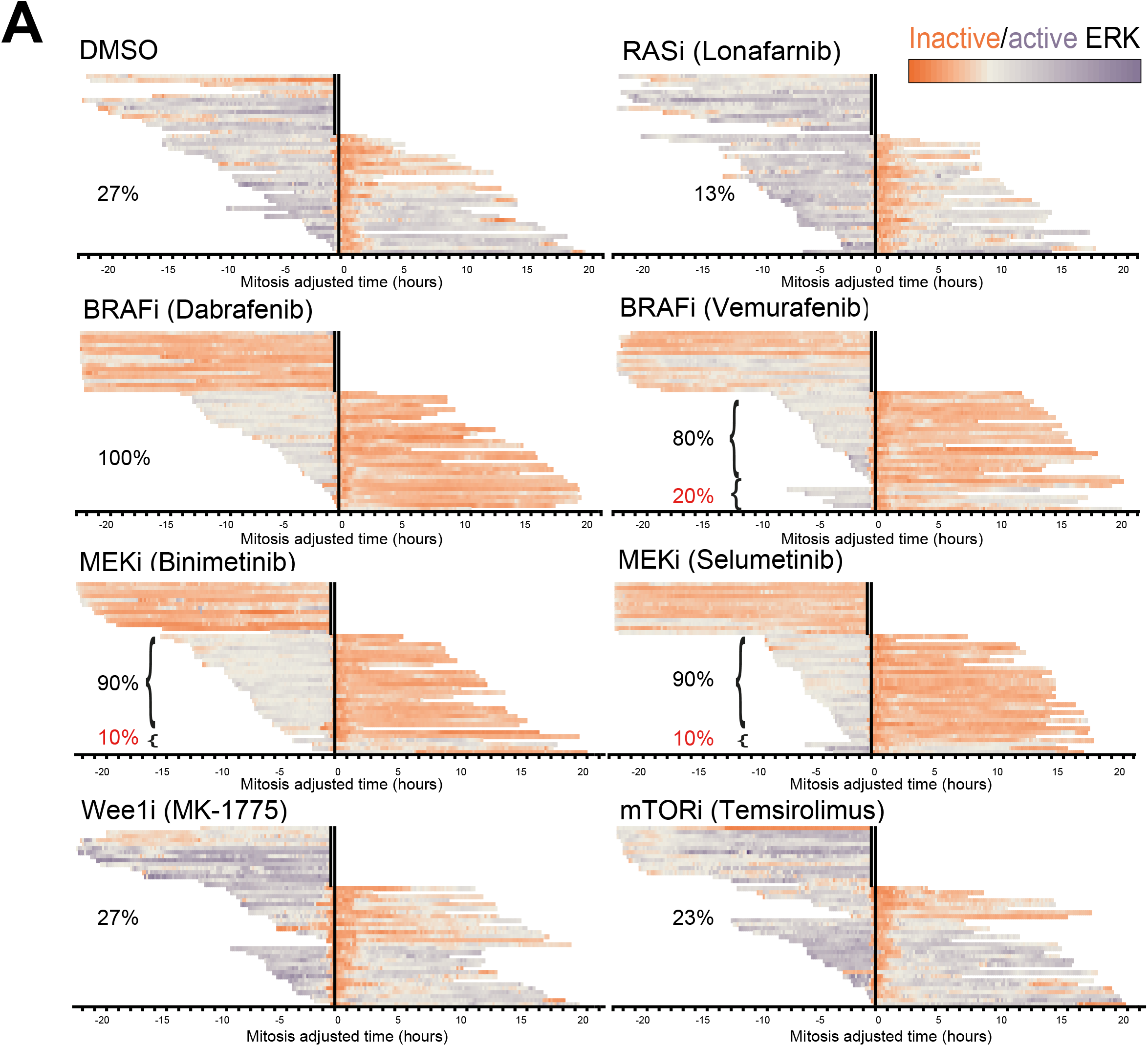
ERK activity in single cells following treatment with inhibitors of BRAF and MEK kinase activity. (A) Each row is an individual cell track of ERKTR activity shown as heat-map (x=0). A total of n=45 cells are shown for each treatment condition, including 30 cells which divided at least once and 15 cells which did not. In black are the percentages of cells where ERKTR falls below a threshold, either prior to- or post-mitosis, at which we consider ERK active. In red are percentages of escapers whose ERK activity never falls. Percentages are calculated from the cells shown above which divided, n=30 in all cases. To qualify as a cell which switched off ERK the average of ERKTR ratio was required to be significantly lower after mitosis than before, as determined by with a minimum length of 12 time-points (2hours).

While the addition of BRAF and both MEK inhibitors at sub-lethal concentrations (80 nM) (Supplementary Figure 5), caused an eventual decrease in ERKTR ratios, the decrease occurred with two different types of kinetics. If BRAF or MEK is inhibited more than 13 hours prior to mitosis, the ERKTR also decreased immediately. Given that the average length of S-phase in DMSO-treated A375p^ERKTR^ cells was 6.5 hours (n=20), and the average G2 length was 4.3 hours (n=13), 13 hours prior to mitosis corresponds to a point in G1. However, if BRAF or MEK activity is inhibited less than 13 hours before mitosis, after G1, the ERKTR ratio remained high, or mildly decreased, until mitosis where it then dropped to a nadir (Figure 3). Inhibition of Wee1 or mTOR has no effect on ERKTR ratios compared to DMSO treated cells. Taken together these observations suggest that BRAF or MEK inhibition results in immediate ERK inactivation if the cells are in G1. If cells are in S or G2, BRAF or MEK inhibition will not result in complete ERK inactivation until the subsequent G1. Thus, the cell cycle stage during which BRAF or MEK is inhibited dictates the effect of this inhibition on ERK activity.

### Inherent resistance to BRAF/MEK inhibition exists in isogenic cell lines

Although in Dabrafenib-treated cells 100% of tracked cells eventually inactivated ERK, in Vemurafenib-treated cells, 20% of cells never inactivated ERK (Figure 3A). For both MEK inhibitors, 10% of cells did not inactivate ERK. This suggests that a subset of cells, even in a clonal cell line, are able to escape from the effects of BRAF and MEK inhibition altogether, but that BRAF inhibitors differ in their ability to suppress ERK activity in these cells.

### Melanoma cells enter a quiescent-like state despite BRAF mutation

We hypothesized that the inactivation of ERK in G1 drives cells into an extended G1 arrest; which we defined as a cell not entering S-phase for 600 minutes or more (56). When treated with DMSO, 18.2% of cells divided twice, 36.4% entered s-phase, and 45.5% entered G1 arrest (n=22). Of all cells treated with MEK or BRAF inhibitors (n=104): 26.9% cells entered S-phase, and 11.1% of these cells underwent a subsequent mitosis; but 73.1% of cells did not enter S phase. Of the cells that inactivated ERK post-mitosis, no cells divided a second time during the remaining imaging time (up to 21 hours), whilst four of the cells with active ERK following mitosis divided a second time during the experiment (Table 1.).

By measuring phospho-RB levels in A375p^ERKTR^ cells following 24 hours treatment with DMSO, Dabrafenib and Binimetinib we confirmed that a fraction of melanoma cells enters a quiescent-like state at every mitosis, and that BRAF and MEK inhibition drives cells into this state.

### ERK activity is less sensitive to BRAF/MEK inhibition in S or G2 versus G1

To precisely determine how cell cycle stage affected the ability of BRAF or MEK inhibitors to decrease ERK activity. We imaged A375p^ERKTR/PCNA^ cells for 13.5 hours *before* adding either BRAF inhibitor Dabrafenib of MEK inhibitor Binimetinib. Consistent with our observations of cells that were treated with SMIs prior to imaging (Figure 3), we observed a large decrease in ERKTR ratio immediately in cells that were in early G1 when both drugs were added, with a few notable exceptions added (Figure 4A, B). In cells that were in G1 upon inhibitor addition, but entered S-phase shortly afterwards (late G1 cells), ERKTR ratios did not fall immediately, but instead remained high until the subsequent G1. Cells that were in S-phase or G2 at the time the inhibitors were added, showed a small decrease in ERKTR ratios, but the ratios remained relatively high levels until the next M/G1, whereupon the ratios dropped to their nadir (Figure 4B,C). The proportional decrease in ERKTR following SMI addition in S-phase or G2, vs the decrease following mitosis, was 30% to 60%. To confirm that ERKTR ratios were less sensitive to BRAF or MEK inhibition in G2 and S phases versus G1 phase, we stained fixed A375p^ERKTR/PCNA^ cells for CyclinA2, which is only expressed in S-phase and G2 and classified cells as being in different cycle phases using custom image analysis (Figure 4D, Methods). G1 cells showed a significant decrease in ERKTR ratio (p>0.001) after a 2 hour drug treatment while G2 and S-phase cells still have a significant (p>0.001), but much smaller, decrease in ERKTR ratios. Thus, consistent with our previous observations where SMI were added to asynchronous cultures prior to imaging, inhibition of BRAF or MEK suppresses a fraction of the ERK activity in S/G2, but the maximal effects of this inhibition occurs only following mitosis.

### BRAF-MEK-ERK activity is dependent on cell cycle stage in multiple BRAF mutant melanoma lines

Our observations suggest the existence of a positive feedback loop in A375p cells that is engaged during a point in G1 that can maintain ERK activity even following inhibition of BRAF or MEK. Such inhibition may also repress the actions of a negative feedback loops on ERK that are engaged following G1-S progression. A quiescent-like state must also negatively regulate BRAF-MEK-ERK activity that is otherwise activated by the BRAFV600E mutation. To determine if cell cycle regulation of BRAF-MEK-ERK is a conserved feature of BRAF mutant cells, we first developed an assay where we could rapidly monitor the effects of BRAF or MEK inhibition on G1, S/G2, or quiescent-like cells on ERK activity at a population level. In this assay we treated cells with Binimetinib, Dabrafenib or DMSO for 24 hours and 2 hours, and subsequently measured ERKTR ratios. Based on our imaging of live cells, we predicted that at 24 hours the distribution of ERKTR ratios would have a larger population of low ERK activity cells, as the majority of cells would have passed through at least a single mitosis resulting in maximal ERK inhibition. In contrast, a short treatment would have an intermediate distribution, as cells that had undergone mitosis, or that were in early G1 at the time of SMI treatment, would inactivate BRAF-MEK-ERK signaling, but the ERKTR ratio would remain high if cells were in late G1, S, and G2 cells. Indeed, in cells treated with either Binimetinib and Dabrafenib for 2 hours the shift in distributions of ERKTR ratios were incomplete compared to 24hrs; and in the case of A375p cells bimodal (Figure 5a,b). We transiently transfected the ERKTR into two additional BRAFV600E cell lines, MEL624 (60) and MEL888 (61), and a BRAF V600D cell line WM266.4 (62), and observed 2hour SMI treatment also showed distributions with incomplete inhibition of ERK after 2hours (Figure 5c), thus demonstrating that bifurcation in ERK activity during G1 is present in other BRAF mutant melanoma lines.

**Figure 5.**
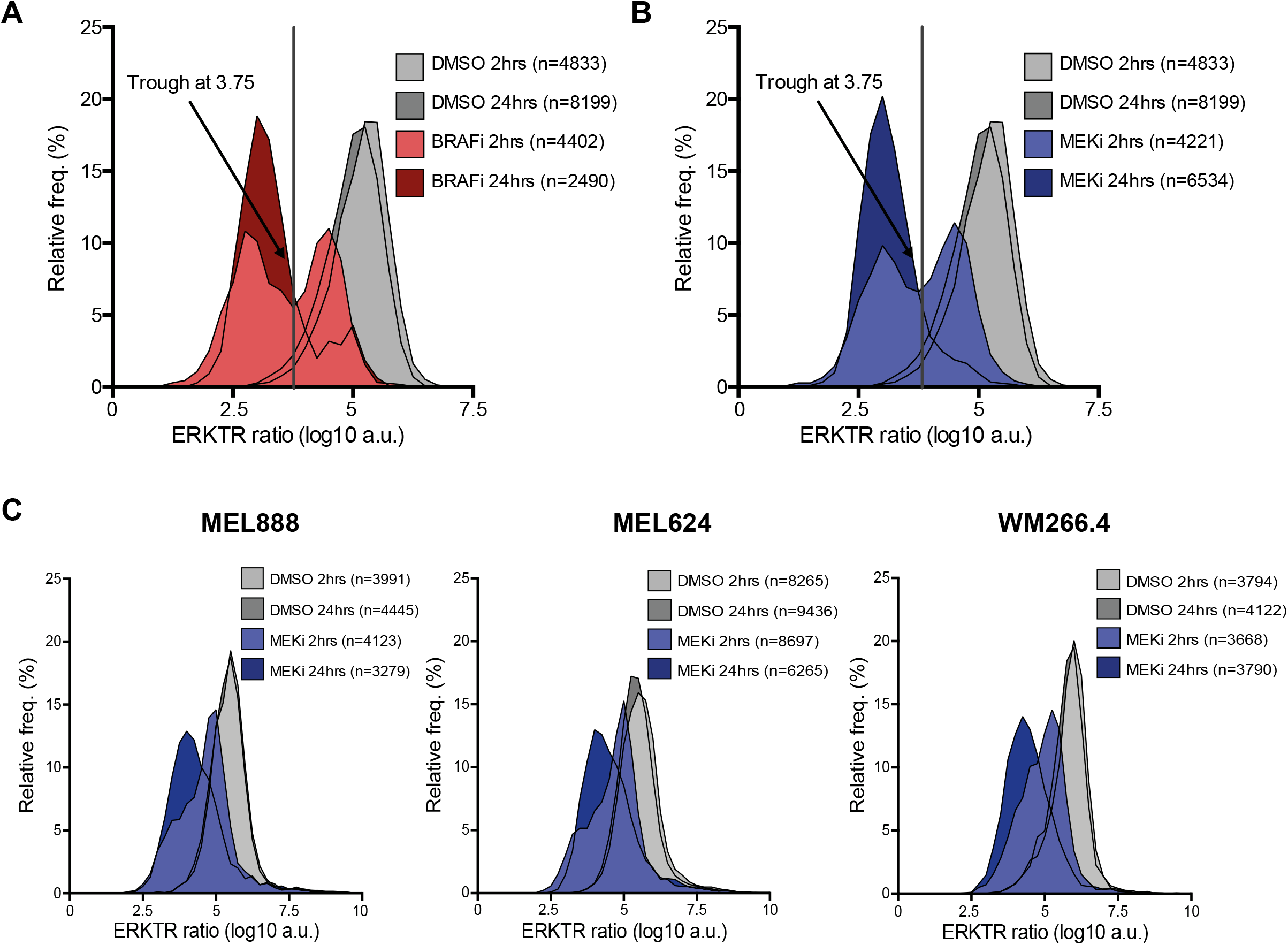
Effect of short term BRAF or MEK inhibition on ERK activity. (A) Distribution of ERKTR ratio following 2 or 24 hours treatment with 80nM Dabrafenib (BRAFi). (B) Distribution of ERKTR ratio following 2 or 24 hours treatment with 80nM Binimetinib (MEKi) or DMSO. (C) ERKTR ratios following 2hours or 24hours with 80nM Binimetinib (MEKi) or DMSO in MEL888, MEL624, or WM266.4 cells.

### Passage through the Restriction Point initiates a positive feedback on ERK

Our live cell data suggested the point where the regulation of ERK activity changes from BRAF-MEK dependent, to independent, is near the Restriction Point (RP). We hypothesized passage through the RP may engage a positive feedback loop that promotes ERK activity even following decreases in BRAF and MEK activation. ERK itself drives passage through the RB by upregulating levels of Cyclin D1, which activates CDK4/6 phosphorylation and concomitant phosphorylation of the RB pocket proteins (68). To determine how ERK dynamics are coupled to RP passage, we quantified ERKTR ratios, and RB phosphorylation (p-RB) simultaneously in single cells following shortterm BRAF or MEK inhibition (2 hour). Using flow cytometry with Hoechst staining for DNA content, we confirmed that in A375p cells p-RB levels increase in G1 and high levels are sustained into S-phase and G2 (Supplementary Figure 6). Untreated A375p cells separate into proliferating ERK^high^/p-RB^high^ cycling population, and an ERK^high^/p-RB^low^ populations, the latter of which is comprised of cells in early G1 cells where ERK activity is rising, but CDK4/6 activity has not yet been maximally activated. A population of ERK^low^ /p-RB^low^ cells also exists consistent with our observations that ERK activity in single cells can bifurcate at mitosis into proliferating and quiescent-like states (Figure 6A). Simultaneous quantification of p-RB and ERKTR localization revealed cell-to-cell variability in p-RB levels in cells with low ERK activity following 2 hour Dabrafenib or Binimetinib treatment; as there is both an ERK^low^/p-RB^low^ state, and an ERK^low^/p-RB^high^ state (Figure 6A, B). These data suggest that while cells pre-RP are largely dependent on BRAF-MEK-ERK signaling for CDK4/6 activity, cells can cross the RP in the absence of ERK activation; presumably once Cyclin D1 levels have sufficiently accumulated to drive progression. In comparison, treatment with a CDK4/6 inhibitor gave a small reduction in ERKTR ratio and a significant change in pRB levels (Figure 6E). CDK2i gave a slight increase in ERKTR ratio and a decrease in pRB (Figure 6D). The proportions of cells in quadrants of pRB and ERK activity under these conditions are shown in Figure 6E and a model of how these states relate to the restriction point is shown in Figure 6F. When taken together with our previous observations that ERK activity can remain high if BRAF and MEK are inhibited after the RP, this suggests that passage through the RP, but not CDK4/6 itself, engages positive feedback loops capable of sustaining ERK independently of BRAF or MEK.

**Figure 6.**
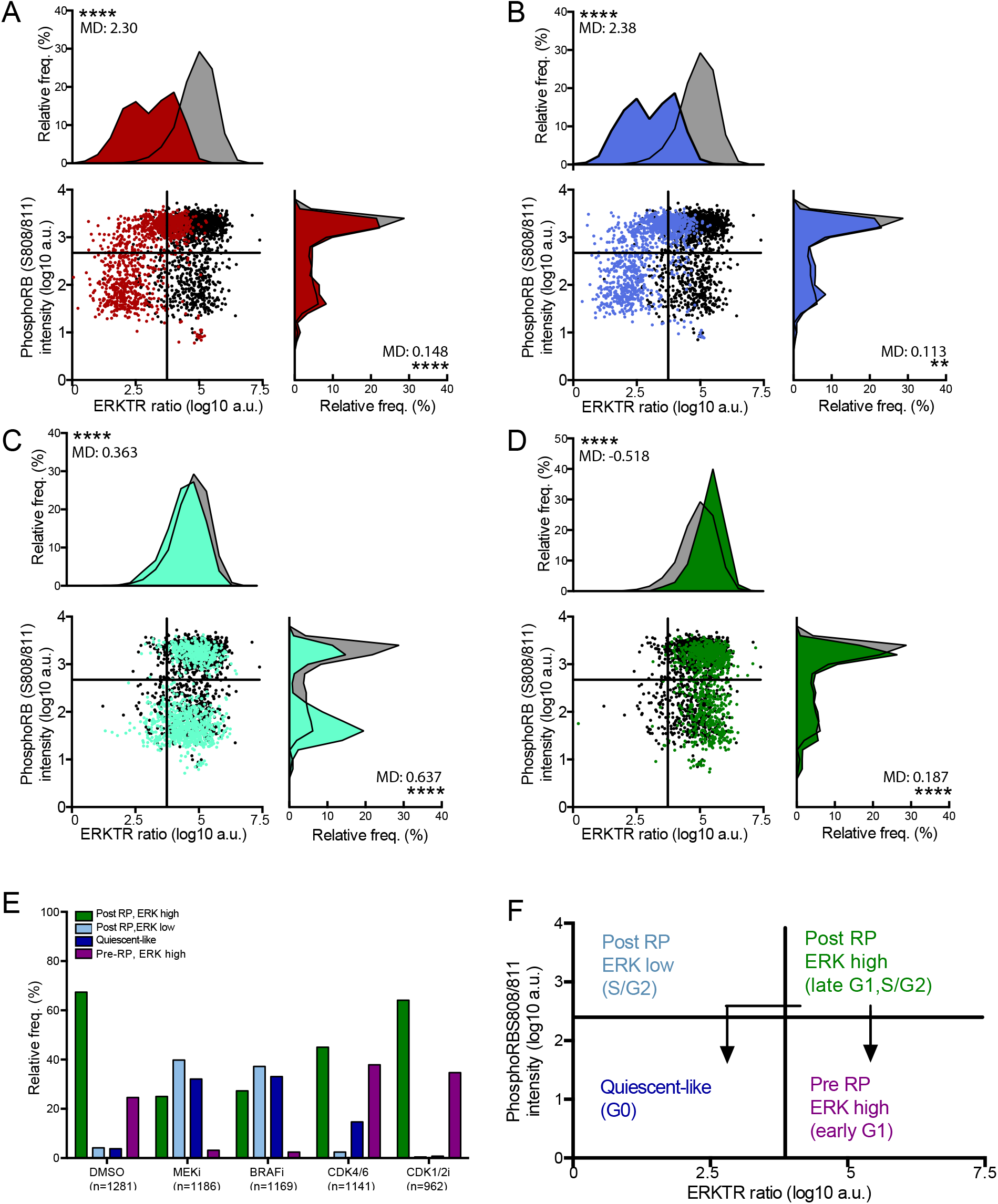
Correlation of ERKTR activity with passage through the RP. (A-D) Scatterplots of ERKTR ratio (x-axes) vs nuclear S808/S811phospho-RB (y-axes). Frequency distributions of ERKTR ratio and phospho-RB intensity are shown above and to right of each scatterplot respectively. Cells were treated for 2 hours with either DMSO or: (A) 80nM Dabrafenib (BRAFi); (B) 80nM Binimetinib (MEKi); (C) 100nM Palbociclib (CDK4/6i); or (D) 100nM CDK1/2 Inhibitor III. (E) Proportions of ERKTR^high^/pRB^high^, ERKTR^low^/pRB^high^, ERKTR^low^/pRB^low^, ERKTR^high^/pRB^low^ in (A-D). (F) Predicted cell cycle stage of cells with different ERKTR/pRB levels

The observation that ERK activity pre-RP, but not post-RP, is dependent on BRAF and MEK opens a therapeutic avenue, as blocking passage through the RP might render cells more sensitive to BRAF or MEK inhibition. To test this idea, we quantified ERKTR ratios and p-RB levels following inhibition of CDK4/6 activity by Palbociclib and/or following treatment of cells with a combination of Palbociclib and BRAF or MEK inhibitors. As expected, treating cells for 2 hours with the CDK4/6 inhibitor Palbociclib increased the fraction of p-RB^low^ cells from 28% to 52%. ERKTR activity was largely not affected by short-term Palbociclib treatment - consistent with the idea that BRAF and MEK promote ERK activity in pre-RP G1 cells. However, combining Palbociclib with BRAF (Figure 7A, C) or MEK (Figure 7B, C) inhibitors in the short-term caused a marked increase in the number of ERK^low^/p-RB^low^ compared to short-term treatment of BRAF, MEK, or CDK4/6 inhibition alone (Figure 7C). Strikingly, the effect of short-term combined CDK4/6 and BRAF/MEK inhibition was even observable on the whole population level, and not just in p-RB high/low populations (Figure 7D). Together these data suggest that Palbociclib sensitizes ERK activity to BRAF or MEK inhibition by preventing passage through the RP and the transition from a BRAF/MEK dependent to independent state.

**Figure 7.**
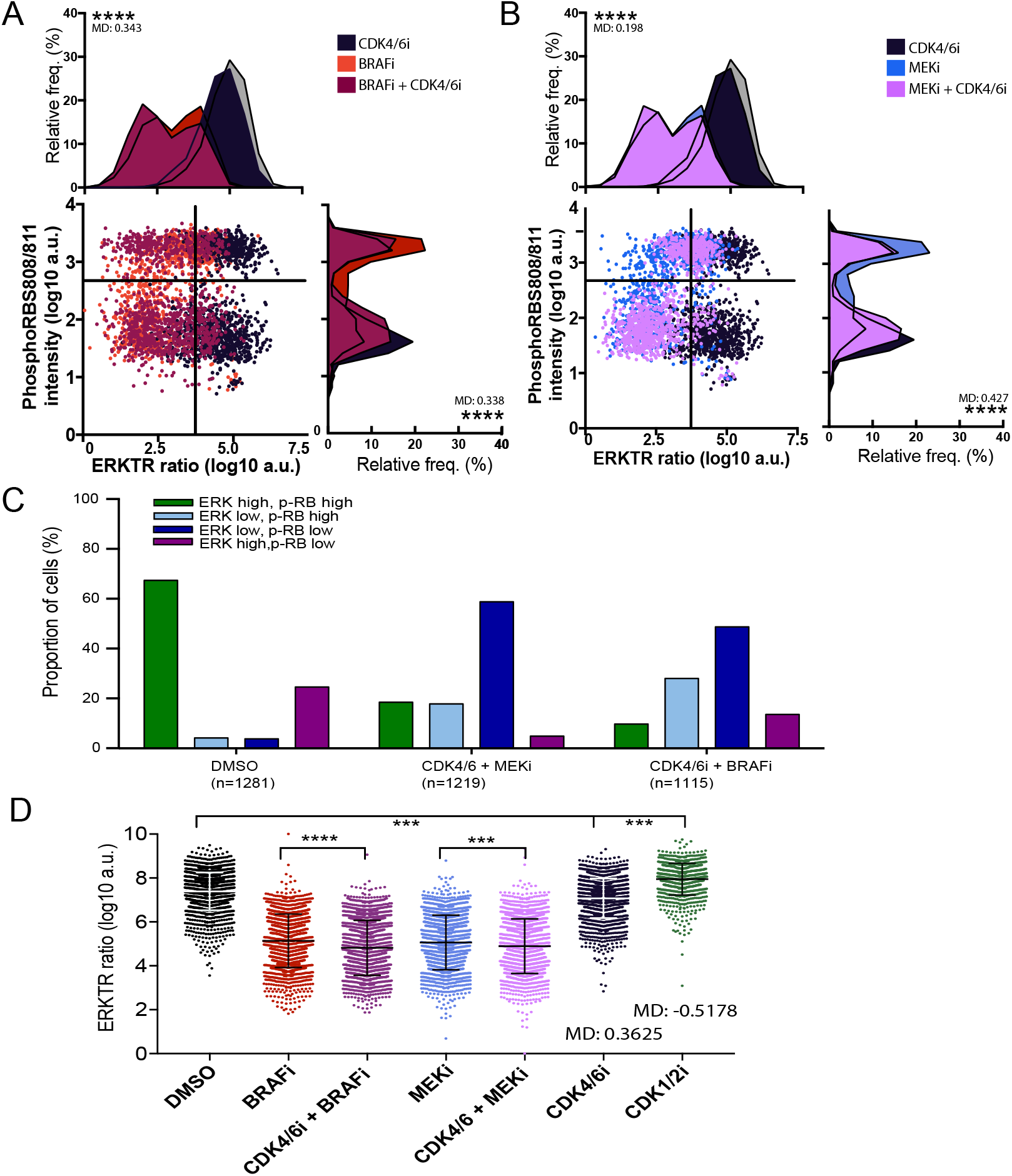
ERKTR ratios after combining CDK4/6 and BRAF or MEK inhibitors. (A) ERKTR ratios and phospho-RB levels after combined treatment of A375p^ERKTR^ cells 80nM Dabrafenib (BRAFi) and 100nM Palbociclib (CDK4/6i). (B) ERKTR ratios and phospho-RB levels after combined treatment of A375p^ERKTR^ cells 80nM Binimetinib (MEKi) and 100nM Palbociclib. (C) Proportions of ERKTR^high^/pRB^high^, ERKTR^low^/pRB^high^, ERKTR^low^/pRB^low^, ERKTR^high^/pRB^low^ cells in (A) and (B). (D) ERKTR ratios in entire populations of single cells from (A) and (B), *** indicates p<0.001; mean differences are shown.

In normal cells, mitogenic signaling converges on CDK2 activity, which together with the extent of DNA damage determines the outcome of the proliferation-quiescence decision (23, 24, 56, 67, 68). We thus sought to determine the role of BRAF-MEK-ERK signaling in the regulation of CDK2 activity following BRAF or MEK inhibition; by monitoring CDK2 activity using a reporter we have developed (66). Notably, short (2 hours) inhibition of either BRAF or MEK had little immediate influence on CDK2 activity (Figure 8A), although a population of ERK^low^/CDK2^low^ cells already exists at 2 hours (Supplementary Figure 7). But CDK2 activity was significantly reduced after 24 hours exposure to BRAF and MEK inhibitors (Figure 8B), as the cells enter a quiescent state-like. This suggests that in melanoma cells, BRAF-MEK-ERK signaling promotes proliferation via upregulation of CDK2 activity, but once CDK2 is activated, BRAF-MEK-ERK activity is dispensable for maintenance of CDK2 activation. This bistable aspect of CDK2 activity that is dependent on a window of mitogen stimulation in normal cells, is thus also an inherent property of cells with activating BRAFV600E mutations.

**Figure 8.**
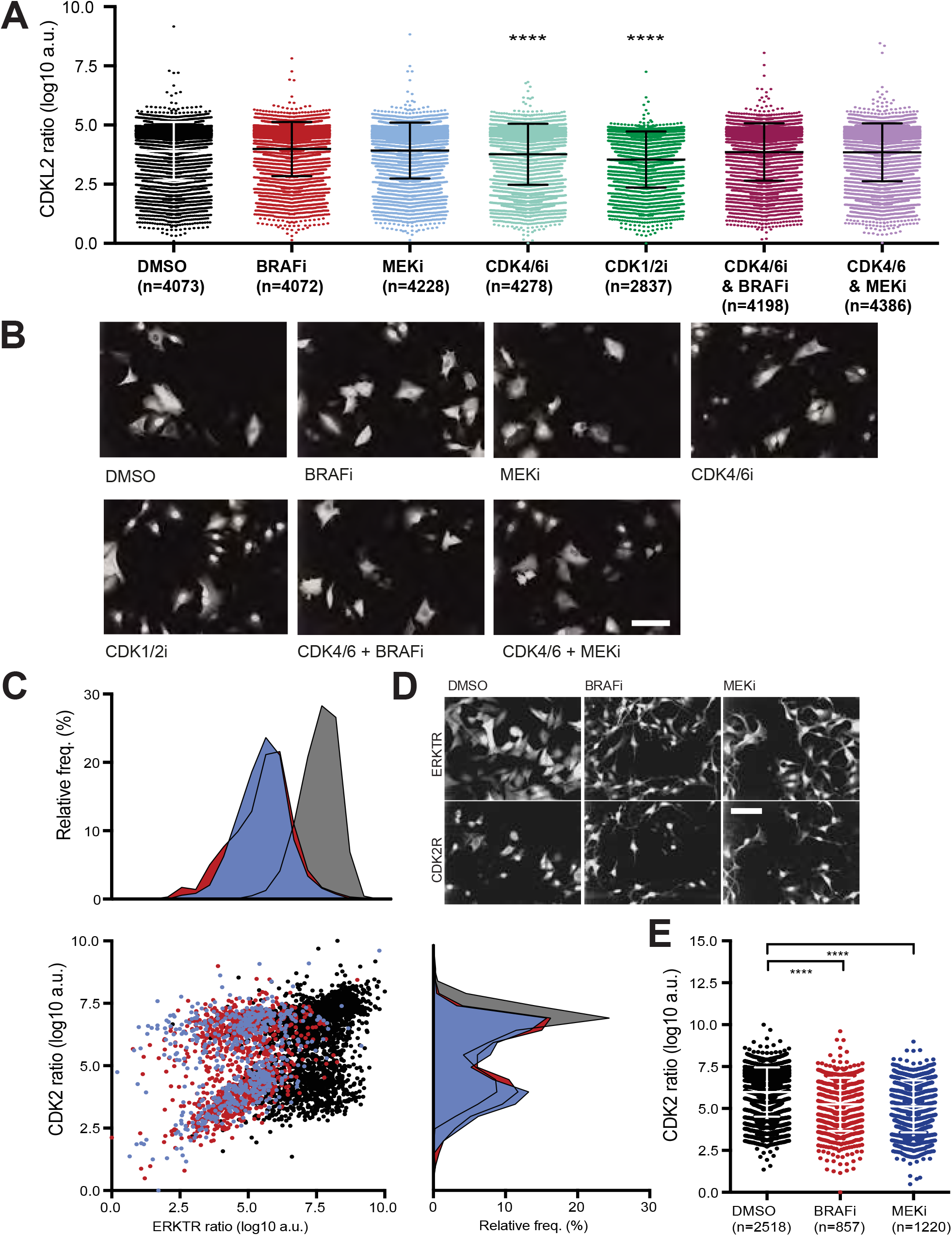
CDK activity in melanoma cells following BRAF and MEK inhibition. (A) CDK2L-GFP ratio (Scaled log10 cytoplasmic/nuclear intensity) after treatment with DMSO, 80nM Dabrafenib, 80nM Binimetinib, 100nM Palbociclib, CDK1/2 inhibitor, combined inhibition with Palbociclib and Dabrafenib, or Palbociclib and Binimetinib and images of cells in different drug treated conditions. (B) ERKTR and CDK2L ratio in single cells in DMSO, Dabrafenib, and Binimetinib treated cells. Images of ERKTR and CDK2L-GFP in single cells, quantification of CDK2L2 ratios after 24hrs. **** indicates significance of p<0.0001, mean differences are indicated. (C) ERKTR ratios of cells treated with CDK inhibitors for 24hrs and MEK inhibitor 2hours. Mean differences shown (n=3 wells). Proportions of cells in each cell cycle phase.

If populations of cells exposed to BRAF or MEK inhibitors for 24 hours are classified by CDK2 activity into CDK2^low^ cells having CDK2L ratio <5 and CDK2^high^ cells having CDK2L ratio >5, there is a correlation between CDK2 activity and ERK activity (R^2^ = 0.37 in MEKi treated cells; R^2^ = 0.45 in BRAFi treated cells) (Supplementary Figure 7b). This suggests that ERK activity is responsive to CDK2 activity, and when CDK2 activity falls below a threshold, ERK activity also drops. We considered that pretreating cells with CDK4/6 and CDK2 inhibitors which induced an increased proportion of cells in G1 would prime cells for MEK inhibition, resulting in a higher than expected mean difference in ERKTR after 2hour treatment of MEKi. In order to test this we treated cells with a selection of CDK inhibitors: CDK1/2 inhibitor 217744, CDK4/6 inhibitor Palbociclib, CDK1 inhibitor R3306, CDK inhibitor Flavopiridol at concentrations which altered the fraction of cell in cell cycle phases after 24hours (Figure 8C). CDK1/2 inhibitor 217744, and CDK4/6 inhibitor showed a greater mean difference in ERKTR ratio when MEKi was added for 2 hours on top than for MEK inhibitor alone (CDK2i 1.42 and CDK4/6 1.12 vs 0.741). This shows that inducing G1 arrest through inhibiting CDK2 or CDK4/6 can prime cells for MEK inhibition and downregulation of ERK signaling, further showing how ERK activity is cell cycle controlled. Conversely R03306 which induced increased fraction of G2 cells, had a lower response to MEKi than the control population (0.519 vs 0.741).

We propose that entry into a quiescent-like state following decreases in ERK activity suppresses the ability of BRAFV600E to activate ERK, establishing a cell-cycle dependent negative-feedback loop.

## Discussion

By quantifying ERK activation dynamics using a reporter in living single cells, we have shown that, despite the presence of an activating BRAF mutation, melanoma populations exhibit considerable inter- and intra-cell variability in ERK activity, both in normal culture conditions and *in vivo*. This observation starkly contrasts with those made largely through bulk analysis methods such as Western blotting which have implied ERK is constitutively active in BRAFV600E cells (3, 71, 72, 73, 74, 74). This variability is primarily due to a bifurcation of cell populations into ERK^high^ and ERK^low^ as cells exit mitosis. Strikingly, clinically relevant inhibitors of BRAF or MEK maximally decrease ERK activity only in G1 cells, but not in S or G2 cells. In S or G2 cells treated with BRAF or MEK inhibitors, the nadir of ERK activity occurs after cells complete mitosis. In both normal and SMI treated cells, drops in CDK2 activity and entry into a quiescent-like state act to inhibit the ability of BRAFV600E to inhibit ERK. Thus, there exists two types of cell cycle regulated feedback on ERK, which affects the response of this signaling pathway to different RAF/MEK inhibitors. In post-RP cells, our data there exists a positive feedback loop which maintains ERK activity in a BRAF/MEK independent fashion, and in quiescent cells ERK activity is suppressed at the levels of BRAF and/or MEK (Figure 9.)

**Figure 9.**
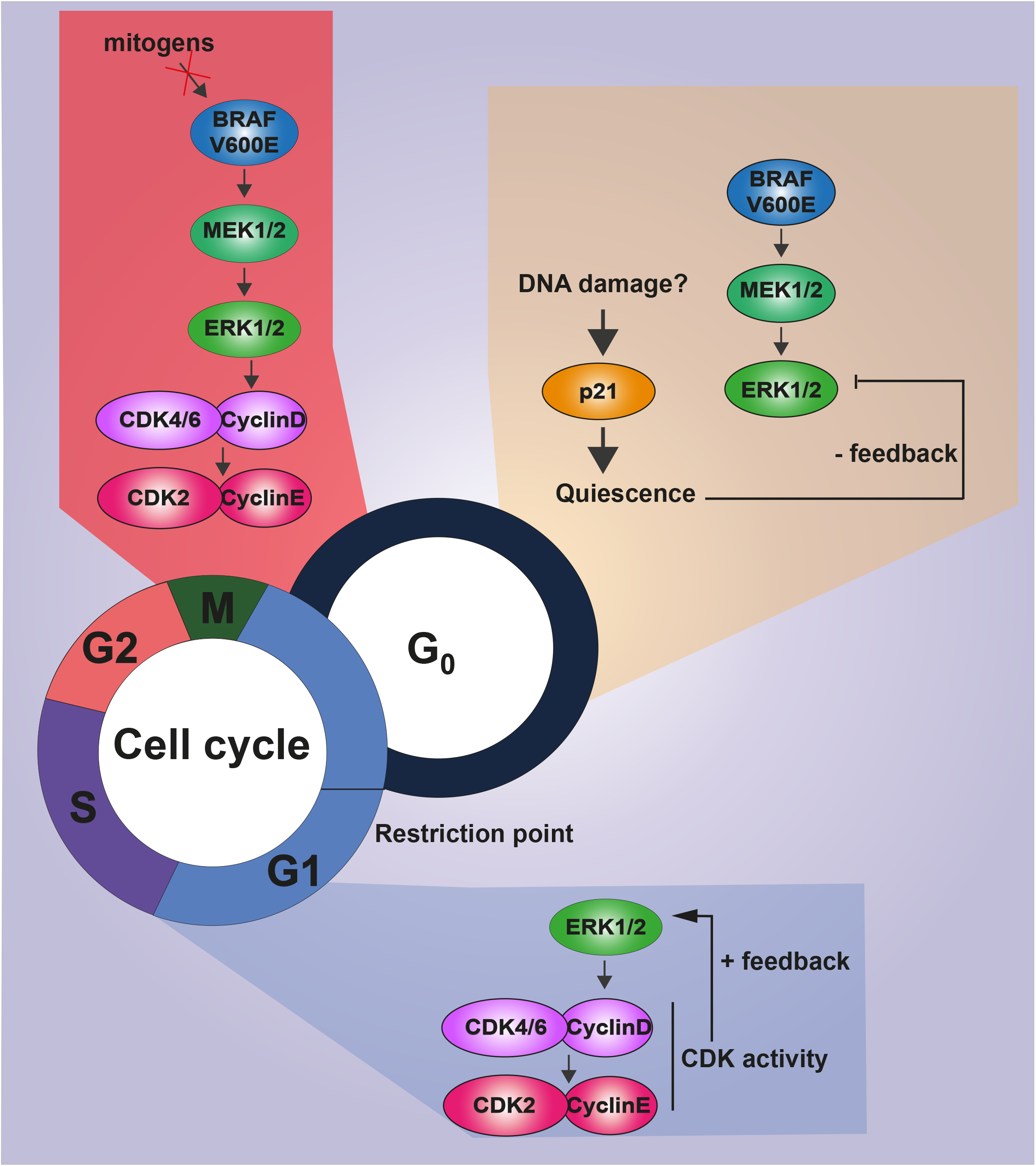
A model for the role of feedback control on ERK activity by the cell cycle. In BRAFV600E cells ERK activity is insensitive to mitogen signaling and promotes G1 entry through upregulation of Cyclin D1. As progression through the Restriction Point (RP), ERK activity is not dependent on BRAF or MEK activity due to positive feedback. In conditions that induce a quiescent-like state, such as endogenous DNA damage and CDK4/6 and CDK2 inhibition, ERK activity is suppressed even in the presence of an activating BRAFV600E mutation.

While the basis for cell-to-cell differences in ERK activity in BRAFV600E cells remains unclear, our data suggests the mechanisms, which promote and/or antagonize BRAF-MEK-ERK activity to generate these differences are controlled in cell-cycle dependent fashions. We propose that, identical to normal cells, the proliferation-quiescence decision in BRAFV600E melanoma cells is still dependent on the mitogenic/pro-proliferative cues via MEK-ERK (23, 24, 67, 68). Whilst the presences of the BRAFV600E mutation largely circumvents the need for exogenous mitogen in melanoma cells, it is insufficient to override the actions of CDK inhibitors (CKIs) such as p21CIP1/WAF1 that can be upregulated following events such an endogenous DNA damage (56, 67, 68). Upregulation of CKIs, and/or down-regulation of CDK activity engages mechanisms that then down-regulate ERK activity (Figure 9). This represents a form of negative feedback control on ERK activity by the cell cycle. That BRAFV600E cells bifurcation into proliferative and quiescent-like states suggests that the ability to enter a quiescent state is beneficial on either a single cell level (i.e. to repair damaged DNA), and/or the population level as it introduces phenotypic heterogeneity into the population which may represent a bet-hedging mechanism against unexpected future stress.

Wild-type BRAF is activated by both recruitment to the membrane by NRAS, as well as by Receptor Tyrosine Kinase (RTK) mediated phosphorylation of residues in the kinase domain (76, 77, 78). Although the V600E mutation can drive full activation of BRAF in melanoma cells that is both independent of RTKs and NRAS (79, 86); some BRAFV600E melanoma cell lines appear to remain dependent on NRAS for proliferation (3) and, BRAFV600E may require NRAS for activation in A375p, MEL888, MEL624 and WM266.4 melanoma cells. Though we do not see a decrease in ERK activation following Lonafarnib treatment, which can theoretically limit NRAS activation by preventing its farnesylation and membrane localization, we note Lonafarnib has little effect on proliferation of NRAS driven melanomas (80, 81); suggesting this compound is not an effective NRAS inhibitor. Alternatively, both wild-type BRAF (82,83,84) and BRAFV600E (85) activity have recently been demonstrated to be upregulated allosterically by KSR-MEK complexes. BRAFV600E-KSR complexes occur at low/intermediate levels of metabolic stress, but at high levels of metabolic stress, AMPK mediated phosphorylation disrupts KSR-BRAFV600E complexes. Thus, KSR and AMPK activity could differ in quiescent versus proliferative cells, leading to differences in the ability of BRAFV600E to activate ERK, and establishing feedback control on ERK by the cell cycle.

Furthermore, mechanisms independent of BRAF-MEK kinase activity must be capable of maintaining ERK activity in S-G2 cells in the presence of BRAF/MEK inhibitors. Our data suggests that these mechanisms are likely to be dependent on increased CDK4/6 and/or CDK2 activity (“CDK activity” in Figure 9). Potentially KSR could act as a scaffold that promotes ERK kinase activity in the absence of MEK, or other ERK kinases could be active S and G2 cells. Conversely, ERK negative regulators such as DUSPs and Sprouty (86,87) could be deactivated to maintain ERK in an active state in the absence of BRAF-MEK activation.

Notably, our single cell analysis reveals there exists a population of cells in which ERK activity never falls following BRAF/MEK inhibition even following mitosis. This implies that in melanoma populations that have pre-existing, versus evolved, resistance to BRAF or MEK inhibitors.

Our results have a number of implications regarding the use of BRAF and MEK inhibitors as treatments for melanoma and possibly other cancers. We show that, as expected, BRAF and MEK inhibitors are effective at inhibiting ERK activity, but only in pre-RP/G1 cells; leading to a quiescent state, negative feedback on ERK, and the stabilization of the ERK^low^ state (Figure 9.). But the ability of S and G2 cells to sustain ERK following passage of the RP in a manner independent of BRAF or MEK could potentially represent a means by which tumor cells could survive exposure to BRAF and/or MEK inhibitors. Moreover, our data suggests that therapeutic combinations that stall cells in mitosis, such as spindle poisons or in G2, such as Flavopiridol (a broad range CDK inhibitor) and RO3306 (a CDK1 inhibitor) may not be able to inhibit ERK successfully. Although we do not propose that simply being in late G1, S, or M is sufficient for cells to survive BRAF or MEK inhibition, we do speculate that differences in cell cycle progression may represent an opportunity for other adaptive or long-term resistant mechanisms to be engaged (8, 89, 90, 91, 92, 93).

Our data also provide rationale for using CDK4/6 inhibitors in combination with BRAF or MEK inhibitors, as the number of cells driven into an ERK^low^ quiescent-like by Dabrafenib or Binimetinib in combination with Palbociclib was greater than following treatment by any single agent. Indeed, such combinations have proven effective in KRAS mutant melanoma, colorectal cancer and Non-small cell lung cancer (94, 95, 96, 97, 98).

A potential caveat to our observations is that the well-established ERKTR reporter (40, 41, 52, 51, 99) may only be detecting a distinct pool of ERK activity. For example, because the reporter detects phosphorylation events at motifs based on the Elk1 protein, it may be may be more sensitive to the activity of ERK molecules, which regulate Elk1 versus those targeting other downstream molecules. Thus, we caution on extrapolating these results broadly to all ERK-regulated events that may occur in melanoma cells. However, because the regulation of Elk1, and other transcription factors represents a key determinant of how ERK drives melanoma progression, our results have significant impact on understanding the role of ERK signaling in tumorigenesis and metastasis; as well as understanding how the NRAS-BRAF-MEK-ERK pathway can be effectively inhibited in patients.

## Methods and Materials

### Generation of fluorescently labeled reporter cell lines

ERK activity was measured using pMSCV-puro-ERKTR-mCherry (donated by Prof. J.G.Albeck). This reporter was originally published in Regot et al, 2014 (40). The average intensities of fluorescence were measured from the nucleus and ring region, and the log10 of the ratio (ring region/nuclear region calculated and referred to as ERKTR ratio. This value was rescaled between 0-10 for clarity in each experiment.

CDK2 activity was measured using pIRES-GFP-PSLD-Puro (CDK2L-GFP) donated by Dr. A.R. Barr. This reporter was originally published in Barr et al, 2016 (66). The average intensities of fluorescence were measured from the nucleus and ring region, and the log10 of the ratio (ring region/nuclear region calculated and referred to as CDK2L ratio).

Cell cycle stage was determined using pIRES-GFP-PCNA-Puro-2b. This plasmid was donated by Dr Alexis Barr, and was originally published in Leonhardt et al 2000 (106).

Monoclonal cell lines were created by transfecting A375p (BRAFV600E) melanoma cells with plasmids using a standard lipofectamine 2000 method. Antibiotic selection was applied for a minimum of 7 days and single cells selected onto a 96well plate by FACS. Cell lines were maintained in DMEM with 10%FBS and 1% Pen/Strep and minimal level of selection agent.

### Live Cell Imaging

Live cell imaging was performed using a High-content Opera Spinning disk confocal microscope (PerkinElmer) with x20 air objective. All live cell imaging was carried out in an environmental chamber at 37°C, 5% CO_2_ and 80% humidity.

Cells were plated 24hours prior to imaging onto 384 well Perkin Elmer Cell Carrier plates in 25ul DMEM (with 10% serum (FBS) and 1% Pen/Strep). SMIs or DMSO were added on top in 25ul at double concentration and plates were centrifuged for 1minute at 1000rpm. Imaging began at 15minutes after drug addition or as specifically stated.

### Feature extraction

After imaging the, files were combined and live cell movies were processed using customized MATLAB scripts for cell tracking and feature extraction as (56) using NucliTrack (58). ERKTR ratio was used to measure ERK activity. The mean intensity of ERKTR fluorescence in a ring region was measured by marking a mask overlaid onto segmented nuclei and being defined as an area of 7pixels wide surrounding the nucleus. The nuclear intensity of EKRTR was measured by taking the average intensity of ERKTR fluorescence in the segmented nucleus. The ERKTR ratio was calculated by dividing the ring region intensity by the nuclear intensity and the Log10 of this was used as the output of ERKTR ratio. Taking the Log10 value of the C:N ratio overcomes the non-equivalence of opposite ratios, briefly if the nuclear fluorescence has a value of 2, and the cytoplasmic has a value of 1, the C/N will be 0.5. Conversely if the nuclear fluorescence is 1 and the cytoplasmic is 2, the C/N ratio would be 2. So even though the ratios are 1:2 and 2:1 (equivalent and opposite) the C/N is not equivalent. Log normalizing these values gives -0.3 and 0.3 respectively, meaning that values in which the denominator is higher are not underrepresented. CDK2 activity was measured in the same way and referred to as CDK2L ratio. Cell cycle stage was demarked using mean intensity of PCNA, which increases in S-phase, and unidirectional standard deviation of PCNA to measure only bright PCNA foci, which appear in S-phase (57). Mitosis was annotated to the track semi-automatically.

Features extracted are:

1. Distance moved: calculated from the X, Y coordinates of two consecutively tracked points from a single cell.

1. Cell area – area (pixels) of segmented nuclei
2. Major axis length
3. Minor axis length
4. Eccentricity
5. Equivalent diameter: diameter of a circle of area equivalent to the segment.
6. Solidity
7. Perimeter
8. Roundness: defined as 4π x area/perimeter^2^
9. PCNA means intensity
10. PCNA standard deviation of intensity
11. PCNA kurtosis of intensity
12. Foci strength: the floored standard deviation of PCNA intensity. This identifies bright spots, ignoring dark ones.
13. ERKTR standard deviation of intensity
14. ERKTR nuclear intensity
15. ERKTR ring region intensity: the intensity of ERKTR in the ring region defined as an area of 7pixels wide from the edge of the segmented nucleus.
16. Mean silhouette: K-means (K=2) clustering of nuclear segments to identify double nuclear segments.

### High Content Immunofluorescence

Cell were seeded onto Perkin Elmer Cell Carrier 384 well plates and allowed to grow for 24hours under normal cell culture conditions. SMIs were added for the time specified (24hours or 2 hours) and cells were fixed at 37°C for 10minutes with 4% formaldehyde. Cells were permeablized for 15mins with 0.01% Triton1000 in PBS at RT, and blocked with 2%BSA in PBS for 2hours. Primary antibodies were diluted 1/1000 in blocking buffer and incubated at 4°C overnight. Alexa Fluorophore-conjugated secondary antibodies (Invitrogen) diluted 1:1000 in blocking buffer were incubated on cells for 1 hour at RT. Hoechst was diluted 1:1000 in PBS and incubated on cells for 10mins at RT. Primary antibodies used are as follow: phosphoThr202/Tyr204 ERK 42/44 (Cell Signaling), phosphoS808/811 RB (Cell Signaling). Fixed imaged were processed using Columbus (PerkinElmer) for cell segmentation and feature extraction.

### Cell Cycle stage determination in fixed cells

Cell populations were first grouped by CyclinA2 intensity to separate G1 cells from S or G2 cells. We then applied a linear classifier on nuclear PCNA texture features to determine cells with characteristic S-phase PCNA foci or G2 cells with smooth PCNA texture. The linear classifier had a goodness of 3.16 and an offset of -13.86. The features measured for the linear classifier were:

- Nucleus Ratio Width to Length
- Nucleus Length
- Nucleus Width
- Nucleus Roundness
- Nucleus Area
- Nucleus PCNA Gabor Max 2 px w2
- Nucleus PCNA Gabor Min 2 px w2
- Nucleus PCNA Haralick Homogeneity 1 px
- Nucleus PCNA Haralick Sum Variance 1 px
- Nucleus PCNA Haralick Contrast 1 px
- Nucleus PCNA Haralick Correlation 1 px
- Nucleus PCNA SER Dark 0 px
- Nucleus PCNA SER Bright 0 px
- Nucleus PCNA SER Saddle 0 px
- Nucleus PCNA SER Valley 0 px
- Nucleus PCNA SER Ridge 0 px
- Nucleus PCNA SER Edge 0 px
- Nucleus PCNA SER Hole 0 px
- Nucleus PCNA SER Spot 0 px
- Intensity Nucleus PCNA Mean

### Tumour xenografts and intravital in vivo imaging

6-8 week old CD1 nude mice were injected subcutaneously with 1 x 10^6^ A375pERKTR cells (n = 2) in a mixture of 50:50 PBS/matrigel. Intravital imaging using a Leica TCS SP8 microscope was performed when tumours reached 6-8 mm, as described (Wyckoff JB et al. 2006). Briefly, mice were anesthetized and minor surgery was performed to expose the tumour. Tumours were excited with an 880 nm pulsed Ti–Sapphire laser and emitted light acquired at 440 nm (collagen second harmonic generation, SHG). In addition, a mCherry signal (ERKTR) was acquired sequentially using confocal microscopy. During approximately 10-min intervals, 5 to 8 different regions were imaged simultaneously for 2 h for each tumour. In each region, a z-stack of 4-5 images (approximately 50 μm deep on average) was taken at a spacing of 10 μm between images, resulting in a time lapse three-dimensional z series for analysis. Z-stacks and time-points were processed in FIJI (93) using 3D drift correction script (94). Intensities were measured using the oval selection tool to manually select nuclear and cytoplasmic regions and normalized to the average of 30 measures in control areas (where no cells were visible) in each image. Mean intensities of nuclear and cytoplasmic regions in the same cell were used to calculate ERKTR ratio, taking the log10 of the normalized cytoplasmic/nuclear intensity and rescaling to 0-10. Measurements were taken from the first time-point and the first drift adjusted z-stack of each position in the tumor. We measured ratios in two camera positions of two biologically independent tumors (the same cell line injected into two separate mice).

### Inhibitors

Inhibitors used were: Vemurafenib (SelleckChem), Dabrafenib (SelleckChem), Trametinib (SelleckChem), Binimetinib (SelleckChem), Selumetinib (SelleckChem), PD0325901 (Tocris), U0126-EtOH (SelleckChem), Temsirolimus (SelleckChem), Lonafarnib (SelleckChem), Salirasib (SelleckChem), MK-1775 (SelleckChem), PD166285 (Tocris), SCH772984 (SelleckChem), Palbociclib (SelleckChem), Flavopiridol (SelleckChem), RO3006 (Merck Millipore), 217714 (Merck Millipore).

## Supplementary Material

### Supplementary figures

Supplementary Fig.1. A375p melanoma cells are BRAF mutant cell line with a mutation at 1799T>A (V600E).

Supplementary Fig.2. ERKTR and phosphoERK respond to BRAF and MEK inhibitors in A375p cells.

Supplementary Fig.3. (Expanded from Fig.1c) ERKTR demonstrates how drugs targeting ERK signaling reduce the activity of ERK after 24hours.

Supplementary Fig.4. Concentrations of SMIs used in live imaging (Fig.3) are sub lethal.

Supplementary Fig.5. ERK dynamics are not randomly distributed across the cell cycle.

Supplementary Fig.6. phospho(S807/811)-RB arises in G1 in A375p cells.

Supplementary Fig.7. CDK2 activity is not significantly reduced after 2hours BRAF or MEK inhibition.

### Supplementary Movies

1. A375p^ERKTR^ cells in DMSO
2. A375p^ERKTR^ cells in RASi (Lonafarnib)
3. A375p^ERKTR^ cells in BRAF-V600i (Dabrafenib)
4. A375p^ERKTR^ cells in BRAF-V600i (Vemurafenib)
5. A375p^ERKTR^ cells in MEKi (Binimetinib)
6. A375p^ERKTR^ cells in MEKi (Selumetinib)
7. A375p^ERKTR^ cells in Wee1i (MK-1775)
8. A375p^ERKTR^ cells in mTORi (Temsirolimus)

## Figure Legends

Figure 4. The effect of MEK and BRAF inhibitors on ERK activation is dependent on cell cycle stage. (A) Traces of ERKTR ratios before and after Dabrafenib addition (80nM, 13.5 hours after imaging begins) color-coded for cell cycle stage during time of Dabrafenib addition. i) Cells early in G1 (navy; n=17) at time of Dabrafenib addition, cells late in G1 (green; n=6); ii) ERKTR ratios after Dabrafenib addition in S-phase decrease 1.4X fold at mitosis compared to the decrease observed immediately in S-phase (purple;n=8); iii) ERKTR ratios after Dabrafenib addition in G2 decrease 2X fold at mitosis compared to the decrease observed immediately in G2 (red; n=8) (B) Traces of ERKTR ratios before and after Binimetinib (80nM, 13.5 hours after imaging begins) addition color-coded for cell cycle stage during time of Binimetinib addition. i) G1 cells (navy; n=27) and late G1 cells (green; n=4). ii) ERKTR ratios after Binimetinib addition decrease 2.6X fold more at mitosis compared to the decrease observed immediately in S-phase (purple; n=2) iii) ERKTR ratios after Binimetinib addition decrease 1.6X fold more at mitosis compared immediately to the decrease observed G2 (n=9). (C) Images and details of a linear classifier to segment fixed cells by cell cycle stage. (D) ERKTR ratios in cells at different cell cycle stages following 2 hours of Dabrafenib or Binimetinib treatment.

## Acknowledgments

The authors thank Dr Alexis Barr (Dynamical Cell Systems team, ICR, London, UK) for the CDK2L plasmid, the PCNA plasmid and for critical reading of the manuscript, Prof. John Albeck (UC Davis, California, USA) for the ERKTR plasmid. We also thank Dr Caroline Springer (Cancer Research UK Manchester Institute, Manchester, UK) and Jordan Holt (Cell Division team, ICR, London, UK) for critical reading of the manuscript. This research was funded by Justin and Lucy Bull, the Stand Up to Cancer campaign for Cancer Research UK, and by Cancer Research UK Programme Foundation Award to C.B. (C37275/ A20146). Nicola Ferrari and Fernando Calvo were funded by Worldwide Cancer Research (Grant 15-0273) and the Institute of Cancer Research. Fernando Calvo is also funded by Cancer Research UK (C57744/A22057) and the Ramon y Cajal Research Program (MINECO, RYC-2016-20352).

